# Differential roles of diet on development and spinal cord regeneration in larval zebrafish

**DOI:** 10.1101/2023.06.20.545707

**Authors:** Emily J. Purifoy, Karen Mruk

## Abstract

The zebrafish is a powerful model organism for studying development and regeneration. However, there is a lack of a standardized reference diet. Most studies evaluate the rate of growth, survival and fecundity. In this study, we compare three diets and their effects on growth and regeneration after a spinal cord injury (SCI). Fish were fed daily for one week with daily measurements of overall length and width of spinal injury. Significant different in length after the trial were observed between live feed and commercial feeds. Similarly, differences in rate of regeneration were observed. Our data highlights the need for establishing a standardized diet for regeneration studies to improve research reproducibility.

## Introduction

Zebrafish are a powerful model organism for studying development and regeneration. One of the limitations with use of zebrafish as a model is the lack of husbandry standardization, particularly for larvae, which are extensively used for developmental and regenerative studies. Diet and nutrition are variables which have potentially profound effects in multiple animal models (1–3), highlighting the need to control for diet within experiments.

Diet standardization in zebrafish laboratories is often an amalgam of research-based guidelines and oldwives tales. Many different diets are available for use in zebrafish including live feeds such as paramecia, rotifers and artemia, (4–6) or commercially formulated diets like powders or flakes. (7, 8) Although formulated diets offer some level of control over the nutritional content, the nutritional content of live feeds is dependent on what they are fed, which ultimately affects what is available to larvae. (9, 10)

Diet impacts many aspects of zebrafish physiology including: adult size and weight, (11) cardiac structures, (12, 13) behavior, (14) as well as reproductive performance. (15–17) Diet has been found to play a significant role in growth up to 15 weeks of age, with an early rapid growth period. (18, 19) Recent studies quantifying skeletal abnormalities up to 20 days post-fertilization (dpf) found dry feed increased caudal-peduncle scoliosis and gill-cover wrinkling (10). In addition, providing high amounts of food, lead to a faster time to reproduction and maturation at the cost of a shortened window for embryo production. (20) However, larval growth at younger ages is not as well quantified.

Despite the numerous options for rearing larval zebrafish, there is a dearth of studies to determine whether different feeds may affect experiments, particularly when using larval zebrafish. We sought to determine whether diet affected early larval development and regeneration after a spinal cord injury. We observed differences in larval growth and survival based on diet. We also observed differences in glial bridging and swim behavior after injury. Taken together, our data suggests that diet and nutrition plays should be considered when designing larval zebrafish experiments.

## Materials and Methods

### Zebrafish husbandry

Adult zebrafish [wild-type AB strain and *Tg(GFAP:EGFP)*] 3-18 months were obtained from the Zebrafish International Resource Center (ZIRC). Adults were maintained at 28.5°C in a recirculating system (iWaki Aquatic) on a 14:10 hr light:dark cycle and fed in the morning with Ziegler’s adult zebrafish diet and in the afternoon with brine shrimp. Water quality of system water is reported in Supp. Tables 1-2. Embryos were obtained through natural outcross matings. Embryos were cultured at 28–30°C in E3 medium containing 5 mM NaCl, 0.17 mM KCl, 0.33 mM CaCl2, and 0.33 mM MgSO4. Embryos were staged as described previously. (21) All animal procedures were approved by the IACUC committee at the University of Wyoming.

### Larvae diet manipulations

Rotifers (*Brachionus plicatilis*; L-type rotifer) were maintained as a continuous culture at a density of 50-200 rotifers/mL in 15 ppt marine salt (Reed Mariculture). Rotifers were fed morning and afternoon with RGComplete (Reed Mariculture). RG Complete contains a minimum of 20% microalgae with a guaranteed analysis including: 2.7% crude protein (min), 0.9% crude fat (min), 0.2% crude fiber (max), 89% moisture (max), and total high unsaturated fatty acids (HUFAs) 64.0 mg/g d.w. (min). Rotifers were harvested fresh for each day of the experiment and the salt was diluted to 5 ppt. Two mL of the diluted culture was then added to the E2 recovery buffer.

Ziegler’s AP100 larval diet (Pentair Aquatic Habitats) was added directly to E2 buffer to a final concentration (10 mg/100 mL). Guaranteed analysis of the AP100 larval diet includes: 50% crude protein (min), 12% crude fat (min), 2.5% crude fiber (max), 10% moisture (max), 15% ash (max), and 1.3% phosphorous. E2 buffer was replaced daily and supplemented with fresh feed. Given the high level of growth in the media, the buffer was supplemented with 0.00003% methylene blue. (22)

Nutritive media developed for mouthless *Xenopus* tadpoles was used as a third diet. (23, 24) The final working recovery buffer was 9.5% Ham’s Nutrient Mixture F12 (Sigma 51651C), 0.5% calf serum (Sigma C8056) in buffer supplemented with methylene blue.

### Water quality

For all diets, zebrafish were fed for 5 days, solutions filtered, and sent to the Analytical Services Laboratory at the Wyoming Department of Agriculture. System water for the adult zebrafish housing system was tested at the same time. Results are reported in Supp. Tables 1 and 2.

### Zebrafish transection

Larvae were transferred to E2 buffer (25) supplemented with penicillin/streptomycin (Pen/Strep, Gibco #15140-122) on the day of transection. Larvae were mounted on flat slides in 2% LMP agarose. The spinal cord was transected with a microblade (WPI #501731) at the level of the cloaca. Larvae were allowed to recover for approximately 2 hours in petri dishes filled with E2 buffer supplemented with Pen/Strep before mounting and imaging. For multi-day experiments, E2 buffer was replaced daily. Larvae were not fed on the day of transection.

### Imaging

Live larvae were mounted flat on a glass slide using 1.5% low melting point agarose, then anaesthetized using Tricane. Larvae were imaged on an Olympus SZX16 stereoscope equipped with an Olympus DP80 camera and SDF PLANO 1XPF objective. CellSens software was used to capture images and saved as TIFFs. Survival was also noted on a daily basis. TIFFs were opened in FIJI ImageJ. To measure body length the segmented line tool was used. The start of the line was at the furthest rostral point of the larvae, the segment was placed just above the cloaca, and the terminal end of the line was placed on the end of the pigmentation at the caudal side of the larvae. To determine whether zebrafish recovered from SCI, images were opened in ImageJ and the presence of a glial bridge was confirmed visually by two individual researchers.

### Locomotion behavior and glial bridging

Larvae were transected as outlined above and their behavior was recorded for 5 days. Behavior experiments were executed between 11:00 AM-1:00 PM every day to ensure reproducibility. A clear 24 well plate with 1 mL fresh E2 supplemented with Pen/Strep was used for the experiments, with one larva per well. Larvae were fed respective diets as outlined above. Locomotion was assessed using ZebraBox and its ViewPoint LS Tracking Software (ViewPoint Life Sciences, Lyon, France). No stimuli were applied, and behavior was video recorded and tracked every 10 minutes for 60 minutes. Total swim distance was found by adding the tracked distances together (twitch: 0-2 mm/s, small: 2-4 mm/sec, large: >4 mm/sec). Following the last day of behavior experiments, larvae were imaged as outlined above to record the absence or presence of a glial bridge.

### Statistics

For all zebrafish experiments, at least two breeding tanks, each containing 2 to 4 males and 2 to 4 females from separate stocks, were set up to generate embryos. Embryos from each tank were randomly distributed across tested conditions, and unfertilized and developmentally abnormal embryos were removed prior to transection. All larvae were included in experiments until day of death. Graphs were plotted using GraphPad Prism. Values for individual fish are plotted, and data is presented as the mean ± standard deviation (SD). For statistical testing, each distribution was assessed using the Shapiro-Wilk test to determine normality. Significant differences were determined using either an unpaired Mann-Whitney t-test or a Kruskal-Wallis ANOVA test with a post hoc Dunn’s test. P values of < 0.05 was considered statistically significant.

## Results

### Rotifer diet promotes larval growth

To determine the effects a rotifer diet had on early larval development, we harvested rotifers and fed larvae ∼100 rotifers/mL from five dpf. Representative images of larval development from 5 dpf to 9 dpf show larvae grew in length and width (Fig. 1A). Larvae fed with rotifers for five days demonstrated a 32% increase in length (Fig. 1B). To determine whether rotifers affected larval growth and regeneration after spinal cord injury (SCI), we next transected the transgenic line *Tg(gfap:EGFP).* This line labels radial glial, which form a pro-regenerative glial bridge after SCI. (26–31) After transection, zebrafish exhibited limited growth (Fig. 1C,D) with only a 16% increase in body length during recovery suggesting that resources are prioritized towards regeneration over growth after an injury in larval zebrafish. Consistent with previous studies, (26, 31, 32) larval zebrafish began the formation of a glial bridge 2–3 days post-injury (dpi) with 74% of larvae forming a glial bridge at 5 dpi (Table 1). As growth after transection was less than that seen during normal development, we next tracked survival (Fig. 1E). Ninety-seven percent of developing larvae survived on a rotifer diet, with 84% surviving after spinal cord transection. In summary, rotifers sustained high levels of growth and regeneration after SCI.

**Figure 1.**
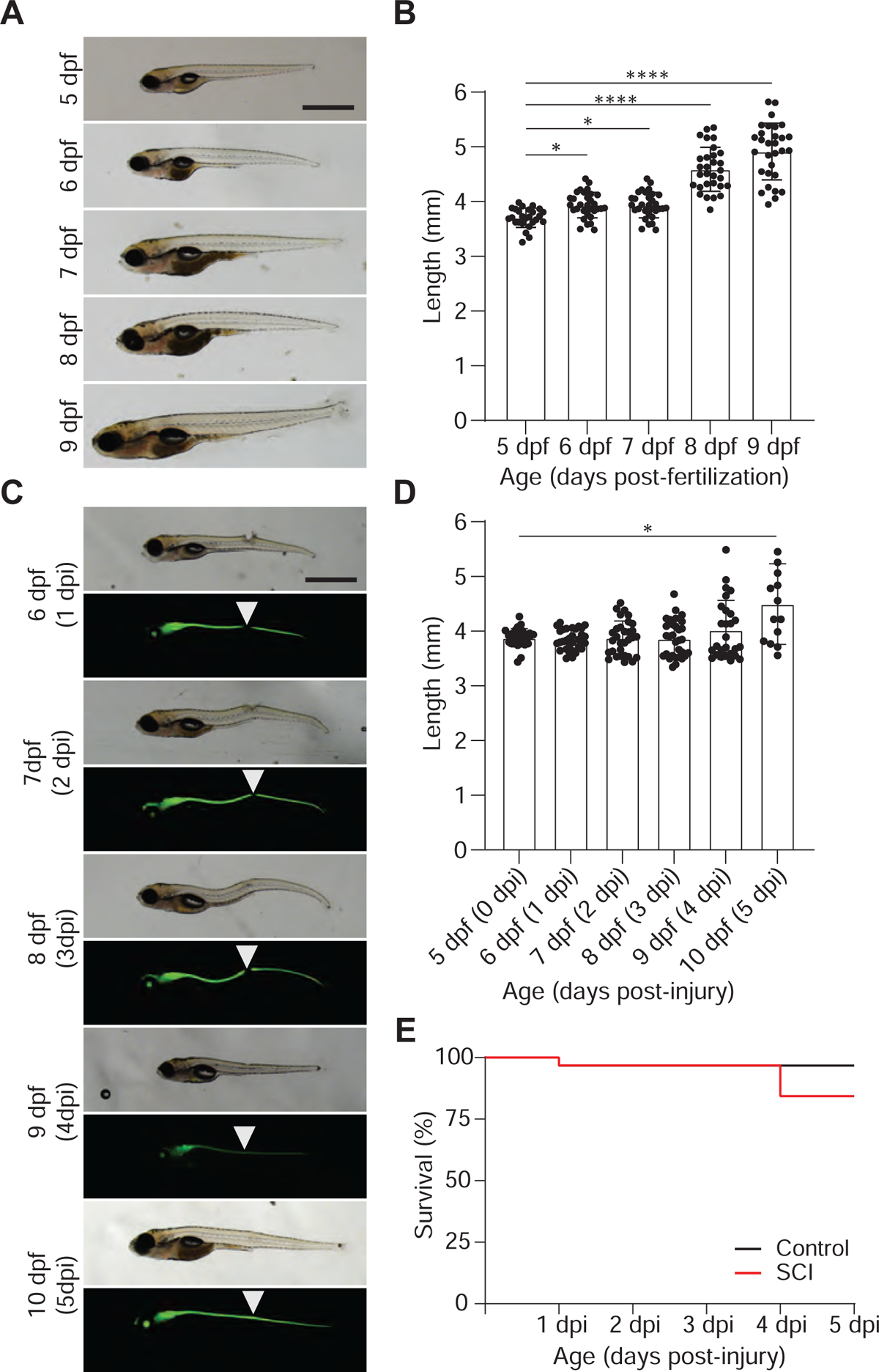
Rotifer-fed larvae have continued growth and regeneration after SCI. (A) Representative bright field micrographs of zebrafish larvae from five days post fertilization (dpf) through 9 dpf are shown. (B) Quantification of larval zebrafish length. Larvae were measured in Fiji and the average length of the larvae ± s.d. plotted. Values for individual fish are shown. The nonparametric Kruskal-Wallis test was used to determine significance. (C) Representative bright field and fluorescent micrographs from *Tg(gfap:EGFP)* larvae are shown after SCI. Arrowhead denotes location of injury. (D) Length of larvae were quantified as in B. (E) Graphs are Kaplan-Meier curves for control (black) and SCI (red). The survival curves for larvae transected at 5 dpf did not significantly differ from control larvae (χ^2^=2.641, p=0.1041). For all micrographs, larvae orientation is lateral view, anterior left. Scale bar: 1 mm. *p≤0.05, ****p≤0.0001

**Table 1.**
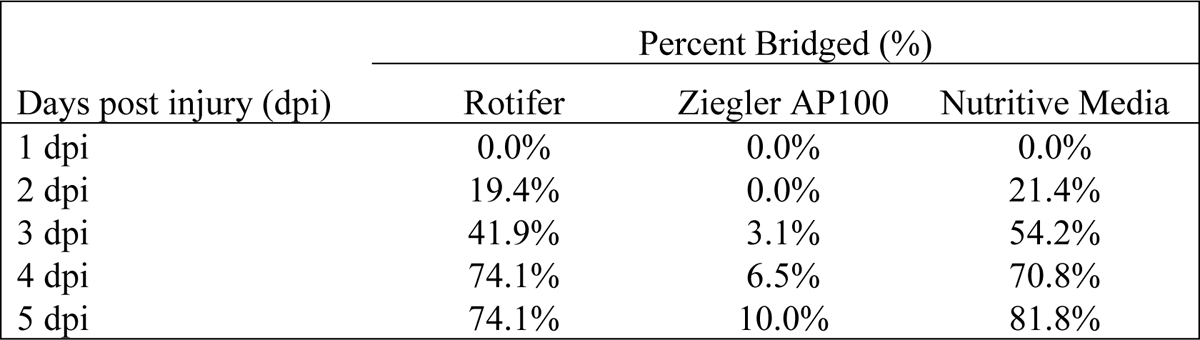
Percentage of larvae forming a glial bridge after spinal cord transection at 5 days post fertilization

By 5 dpf, the nutrition from the yolk sac of zebrafish larvae is mostly exhausted, and larvae begin to rely on exogenous feed sources. (32) We wondered whether larvae injured after yolk sac depletion would be similarly affected by diet. For these experiments, we fed larvae 5-6 dpf, transected fish at 7 dpf, and resumed feeding at 8 dpf. Larvae grew in length until the day of SCI; however, after transection zebrafish larvae showed a small but statistically significant decrease in length with no additional growth (Supp. Fig 1 A,B). Our data suggets after an injury, resources are dedicated to zebrafish regeneration and not developmental growth. We also noticed a significant decline in survival after SCI in larvae injured at 7 dpf and fed a rotifer diet (Supp. Fig. 1C). Of the larvae that did survive, 55% of them were able to form a glial bridge (Table 2). Given the increase in death among injured larvae (<50% survival), we hypothesized that injured fish may struggle to catch the rotifers and sought to increase survival using non-live feeds.

**Table 2.** Percentage of larvae forming a glial bridge after spinal cord transection at 7 days post fertilization

| Days post injury (dpi) | Percent Bridged (%) |  |  |
| --- | --- | --- | --- |
|  | Rotifer | Ziegler AP100 | Nutritive Media |
| 1 dpi | 0.0% | 0.0% | 0.0% |
| 2 dpi | 0.0% | 0.0% | 2.2% |
| 3 dpi | 6.5% | 2.5% | 7.0% |
| 4 dpi | 31.4% | 8.3% | 38.5% |
| 5 dpi | 55.2% | 13.3% | N/A<sup>a</sup> |

<sup>a</sup>All larvae were deceased at time of imaging.

### Powder diet does not promote regeneration from SCI

We first tested Ziegler’s AP100 larval diet as an alternative to live feed. Figure 2A displays representative images of uninjured larvae from 5 dpf to 9 dpf. In contrast to rotifer-fed larvae, zebrafish given the powder diet only grew 4% over the course of the experiment (Fig. 2B). We next transected larvae and found that the powder diet did not sustain continued growth during regeneration, with an overall 7% decrease in length (Figure 2C, D). Moreover, few larvae were able to form a glial bridge (Table 1). Although larvae without injury were found to have a 100% survival over the course of the experiment, injured larvae only had a 60% survival (Fig. 2E).

**Figure 2.**
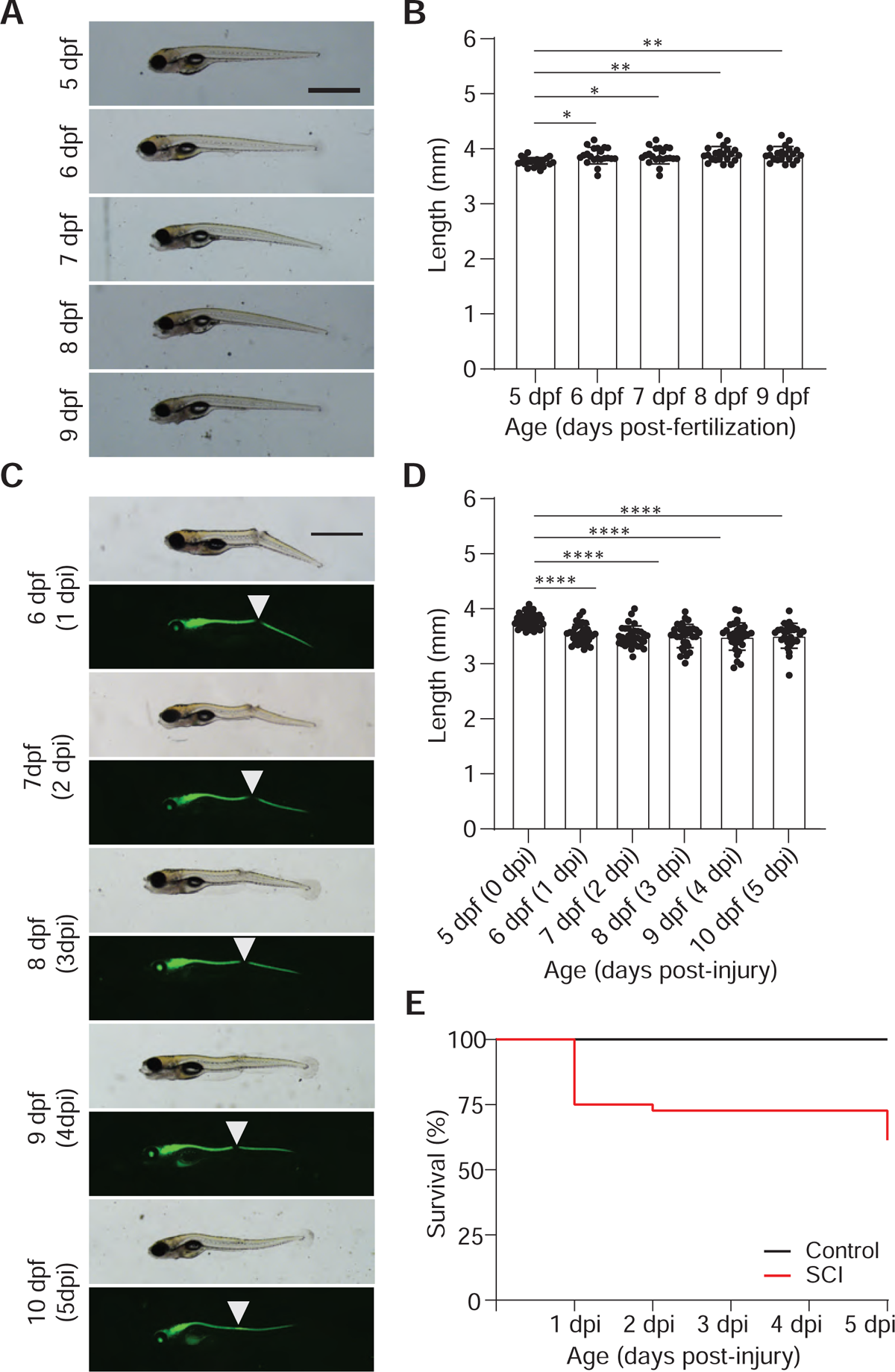
Larvae fed with Zeigler AP100 diet only show continued growth without a SCI. (A) Representative bright field micrographs of zebrafish larvae from five days post fertilization (dpf) through 9 dpf are shown. (B) Quantification of larval zebrafish length. Larvae were measured in Fiji and the average length of the larvae ± s.d. plotted. Values for individual fish are shown. The nonparametric Kruskal-Wallis test was used to determine significance. (C) Representative bright field and fluorescent micrographs from *Tg(gfap:EGFP)* larvae are shown after SCI. Arrowhead denotes location of injury. (D) Length of larvae were quantified as in B. (E) Graphs are Kaplan-Meier curves for control (black) and SCI (red). The survival curves for larvae transected at 5 dpf significantly differed from control larvae (χ^2^=10.41, p=0.0013). For all micrographs, larvae orientation is lateral view, anterior left. Scale bar: 1 mm. *p≤0.05, **p≤0.01, ****p≤0.0001

We also tested the effects of Ziegler’s AP100 on the older larvae transection model (Supp. Fig. 2A). Similar to rotifer-fed larvae, powder-fed zebrafish grew until the day of injury; however, after SCI, demonstrated an overall decrease in length by 1.6%. (Supp. Fig. 2B). Injured larvae given this diet had reduced survival compared to both uninjured and rotifer-fed SCI larvae. (Supp. Fig. 2C). Similar to younger larvae, only 13% of larvae were able to form a glial bridge. As this diet could support development, this data suggests that the resources required for regeneration are different than those required for development.

### Nutritive media does not encourage growth but permits regeneration after SCI

We last tried a nutritive media, as injured fish would not have to seek for or chase food. Representative images of the larvae given this diet from 5 dpf to 9 dpf are shown (Fig. 3A). Uninjured larvae did not grow and saw a decrease in length by 0.5% (Fig. 3B). We next transected larvae and found that the nutritive media did not sustain continued growth (Figure 3 C,D). We observed a decrease in survival for both uninjured and transected larvae (Figure 3E); however, of the transected larvae that survived, 82% of them were able to form a glial bridge by 5 dpi (Table 1) again suggesting that the nutrients required for larval development, survival, and regeneration from SCI are different. This was particularly evident when we transected older larvae (Supp Fig. 3). Larvae transected at 7 dpf and fed nutritive media decreased in length by 9.3% (Supp. Fig. 3B). Nutritive media was unable to support larvae survival as a 100% mortality rate was observed by 5 dpi in both control and injured fish (Supp. Fig. 3C). Surprisingly, 39% of larvae were able to form a glial bridge before death (Table 2).

**Figure 3.**
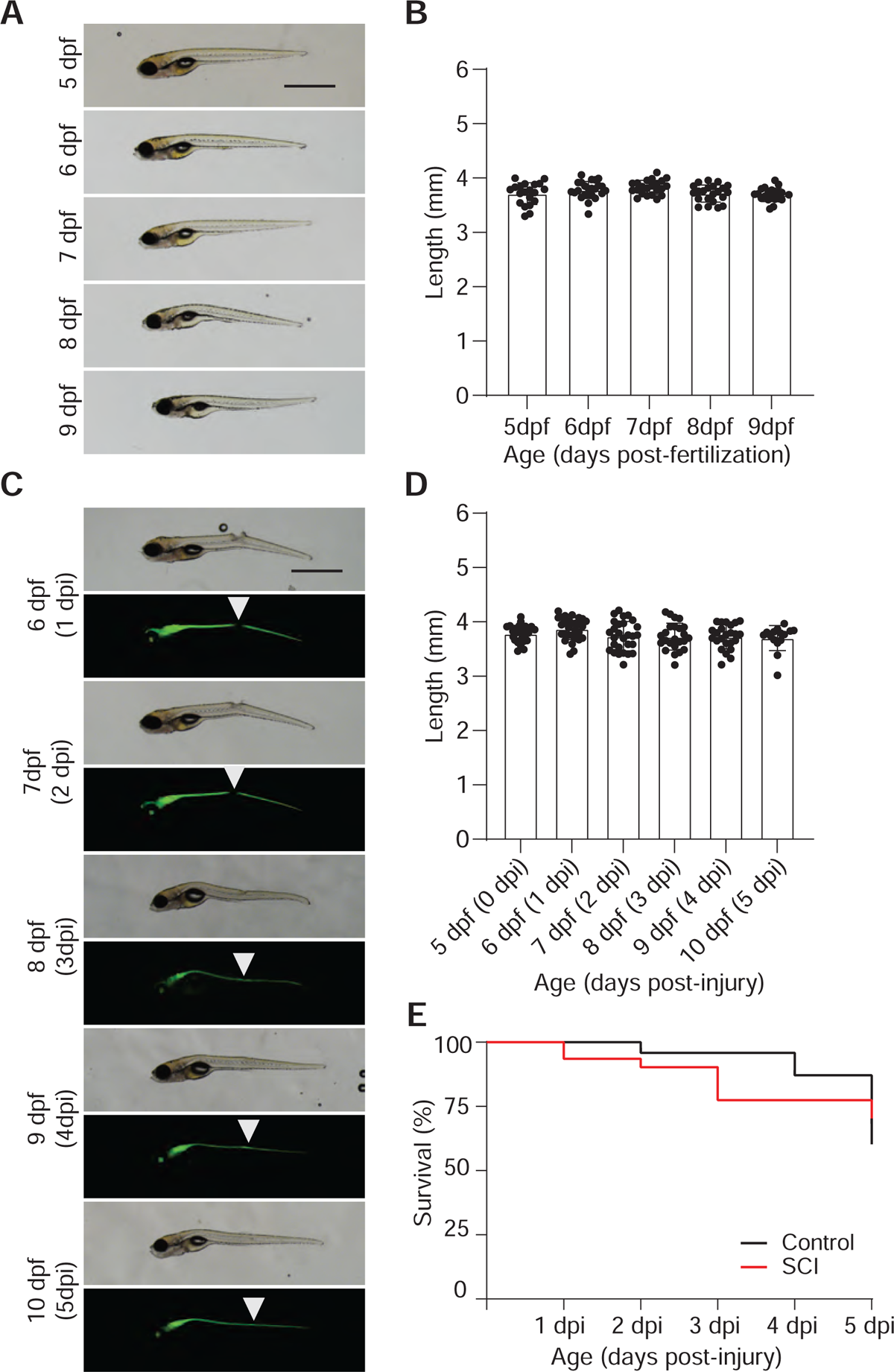
Larvae fed with nutritive media do not exhibit continued growth. (A) Representative bright field micrographs of zebrafish larvae from five days post fertilization (dpf) through 9 dpf are shown. (B) Quantification of larval zebrafish length. Larvae were measured in Fiji and the average length of the larvae ± s.d. plotted. Values for individual fish are shown. (C) Representative bright field and fluorescent micrographs from *Tg(gfap:EGFP)* larvae are shown after SCI. Arrowhead denotes location of injury. (D) Length of larvae were quantified as in B. (E) Graphs are Kaplan-Meier curves for control (black) and SCI (red). The survival curves for larvae transected at 5 dpf did not significantly differ from control larvae (χ^2^=0.0025, p=0.9601). For all micrographs, larvae orientation is lateral view, anterior left.

### Diet affects locomotion in developing and injured larvae

Zebrafish larvae begin freely swimming around 4 dpf. Recent studies have reported changes in swim behavior based on diet; (33, 34) therefore, we chose to look at swimming behavior in uninjured and transected zebrafish larvae. We quantified total swim distances for larvae given each of the diets (Figure 4). In uninjured larvae, overall swim distances decreased as the zebrafish aged with all diets (Figure 4a). After spinal cord transection, we found significant differences in swim distances based on diet. Surprisingly, there was a significant difference in total swim distances as early as the day of injury (Figure 4B). One the day of injury (0 dpi), larvae given the rotifer diet had 143% more swim distance than those given the powder diet, and 257% more than those given the liquid diet. This trend remained throughout the trial, and by 5 dpi, the larvae fed a rotifer diet had total swim distances that were 118% more than those given the powder diet, and 207% more than those given the liquid diet (Fig. 4B).

**Figure 4.**
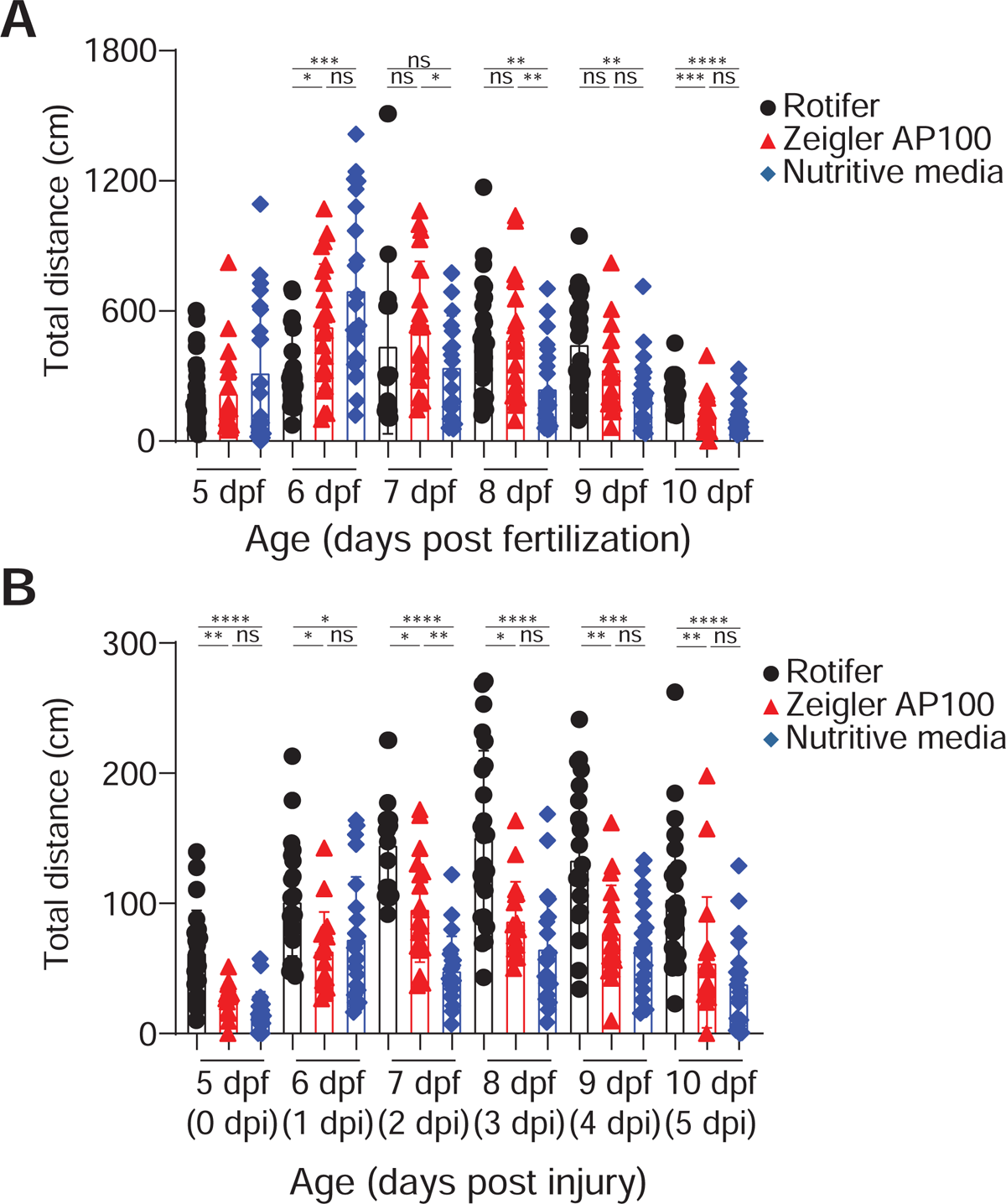
Larvae show differences in total swim distance based on diet. **(A)** WT larvae were followed from 5 dpf through 9 dpf and free swim followed individually. Average swim distances ± sd are shown shown with individual larvae plotted. The nonparametric Kruskal-Wallis test was used to determine significance. (B) *Tg(gfap:EGFP)* larvae were transected at 5 dpf and free swim followed individually. Average swim distances ± sd are shown shown with individual larvae plotted. The nonparametric Kruskal-Wallis test was used to determine significance. Multiple comparisons for each day were compared using a Dunn’s post hoc test. For both panels: *p≤0.05, **p≤0.01, ***p≤0.001, ****p≤0.0001

## Discussion

Diet plays an important role in zebrafish health and reproduction. Despite this, our understanding of how diet effects larval zebrafish experiments remains elusive. To that end, we systematically looked at the role different diets play in development and regeneration after spinal cord transection. Our results demonstrate that different feeds affect both growth and the ability of the larval zebrafish to regenerate from a SCI.

We routinely feed our larvae rotifers to raise lines to adulthood due to the increased survival rate reported. (4, 35) In addition, rotifers are an ideal prey for developing zebrafish larvae because they are smaller than *Artemia*, move slowly through the water column, and promote development of prey capture skills. Because we culture rotifers in nutrient-rich microalgae, supplemented with highly unsaturated fatty acids, it is perhaps unsurprising that larvae fed a rotifer diet grow well.

One unexpected finding was that rotifer-fed larvae and larvae bathed in nutritive media regenerated well after SCI; however, larvae fed the Zeigler larval diet poorly regenerated. A major difference between the rotifer and AP100 diet is the minimum guaranteed crude protein content (2.7% versus 50% respectively). Studies in mammals show that the quantity and source of dietary protein impacts microbiota function. (36, 37) It is possible that similar to mammals, (38–40) bacterial imbalances can exacerbate spinal cord damage and/or impair recovery in zebrafish. The zebrafish community has multiple experimental paradigms to tackle this important question such as: generating genetic mutants in germ-free zebrafish, transplanting exogenous bacteria, and environmental manipulation. (41)

Experimental studies looking at swim behavior in zebrafish have used a range of feeds including: paramecia, (42) *Artemia*, (43) and Zeigler AP100 diet (44) demonstrating the wide variety of feeding protocols used during behavioral experiments. Our study helps bridge the gap in knowledge by studying multiple diets in an experimental context. We found that after a SCI, rotifer-fed larvae had increased swim distances, even with a complete spinal cord transection. One explanation for the observed differences could be explained by an increase in available energy provided from the rotifers. Another explanation is that the swimming behavior of rotifers provides a stimulus to promote swim behavior. Future studies comparing free swimming feeds could provide insight into these differences.

In conclusion, this study highlights that the choice of feed for zebrafish larvae can affect experimental results. Overall, these results have implications for the reproducibility of results across laboratories and highlight the need for publications to provide information on the feed used on experimental animals.

## Supporting information

Supplemental Packet

## Acknowledgements

We thank Thomas Rynes for excellent fish care. We gratefully acknowledge support from the Wyoming Research Scholars program (E.P.).

## Author Contributions

K.M. designed the experimental strategy. E.J.P. completed all of the experiments, data analysis, and figure making. E.J.P. and K.M. wrote and edited the article.

