## Supplemental Packet for "Differential roles of diet on development and spinal cord regeneration in larval zebrafish"

**Table S1. Analytical results of water testing for system water and diets**

| Analysis | Units | System Water | Rotifer Diet | Zeigler Diet | Nutritive Media |
|---|---|---|---|---|---|
| **Alkalinity** | mg/L | <20 | 70.4 | 65.4 | 115.4 |
| **Bicarbonate** | mg/L | <20 | 63 | 65.4 | 107.7 |
| **Calcium** | ppm | 8.72 | 21.41 | 29.22 | 26.7 |
| **Carbonate** | mg/L | <2.0 | 7.4 | <2.0 | 7.7 |
| **Chloride** | mg/L | 270.1 | 704.02 | 672.61 | 978.82 |
| **Conductivity** | µS/cm | 962 | 2680 | 2390 | 3530 |
| **Fluoride**[a] | mg/L | <0.20 | <0.20 | <0.20 | 6.47 |
| **Magnesium** | ppm | 18.64 | 17.34 | 22.28 | 19.28 |
| **Nitrate**[a] | mg/L | 1.77 | <0.20 | <0.20 | <0.20 |
| **Nitrite**[a] | mg/L | <0.20 | <0.20 | <0.20 | <0.20 |
| **pH** | | 7.1 | 8.4 | 8.3 | 8.6 |
| **Potassium** | ppm | 5.85 | 20.9 | 29.69 | 35.97 |
| **Sodium** | ppm | 150.11 | 311.51 | 421.9 | 661.42 |
| **Sulfate+** | mg/L | 41.84 | 110.53 | 107.84 | 126.91 |
| **Total dissolved solids** | mg/L | 503 | 1224 | 1316 | 1916 |
| **Corrosivity** | | <-2.0 | -0.02 | -0.02 | 0.44 |
| **Total Hardness** | mg/L | 98.5 | 124.9 | 164.7 | 146.1 |
| **Nitrate + Nitrite (as N)** | mg/L | <0.20 | <0.20 | <0.20 | <0.20 |

[a] Values are estimated due to the high chloride level in the water

**Table S2. Mineral analysis of water testing for system water and diets**

| Analysis | Units | System Water | Rotifer Diet | Zeigler Diet | Nutritive Media |
|---|---|---|---|---|---|
| Aluminum | ppm | 0.0198 | <0.00899 | 0.0334 | <0.00899 |
| Antimony | ppm | <0.00497 | <0.00497 | <0.00497 | <0.00497 |
| Arsenic | ppm | <0.00689 | <0.00689 | <0.00689 | <0.00689 |
| Barium | ppm | <0.0151 | <0.0151 | <0.0151 | <0.0151 |
| Beryllium | ppm | <0.00301 | <0.00301 | <0.00301 | <0.00301 |
| Boron | ppm | <0.039 | <0.039 | 0.043 | <0.039 |
| Cadmium | ppm | <0.00338 | <0.00338 | <0.00338 | <0.00338 |
| Chromium | ppm | <0.0148 | <0.0148 | <0.0148 | <0.0148 |
| Cobalt | ppm | <0.00498 | <0.00498 | <0.00498 | <0.00498 |
| Copper | ppm | <0.00985 | <0.00985 | 0.0158 | 0.0181 |
| Iron | ppm | <0.079 | <0.079 | <0.079 | <0.079 |
| Lead | ppm | <0.00487 | <0.00487 | <0.00487 | <0.00487 |
| Lithium | ppm | <0.044 | <0.044 | <0.044 | <0.044 |
| Manganese | ppm | <0.00471 | <0.00471 | <0.00471 | <0.00471 |
| Mercury | ppm | <0.000971 | <0.000971 | <0.000971 | <0.000971 |
| Molybdenum | ppm | <0.00500 | <0.00500 | <0.00500 | <0.00500 |
| Nickel | ppm | <0.00477 | <0.00477 | <0.00477 | <0.00477 |
| Platinum | ppm | <0.00499 | <0.00499 | <0.00499 | <0.00499 |
| Selenium | ppm | <0.00491 | <0.00491 | <0.00491 | <0.00491 |
| Silica | ppm | 0.47 | <0.3959 | 2.57 | 0.23 |
| Silicon | ppm | 0.22 | <0.185 | 1.2 | 0.11 |
| Silver | ppm | <0.00493 | <0.00493 | <0.00493 | <0.00493 |
| Strontium | ppm | 0.13 | 0.018 | 0.017 | 0.027 |
| Thalium | ppm | <0.000960 | <0.000960 | <0.000960 | <0.000960 |
| Thorium-232 | ppm | <0.00498 | <0.00498 | <0.00498 | <0.00498 |
| Tin | ppm | <0.123 | <0.123 | <0.123 | 0.027 |
| Titanium | ppm | <0.010 | <0.010 | <0.010 | <0.010 |
| Uranium-238 | ppm | <0.00498 | <0.00498 | <0.00498 | <0.00498 |
| Vanadium | ppm | <0.00478 | <0.00478 | <0.00478 | <0.00478 |
| Zinc | ppm | <0.00469 | 0.00605 | 0.00907 | 0.0334 |

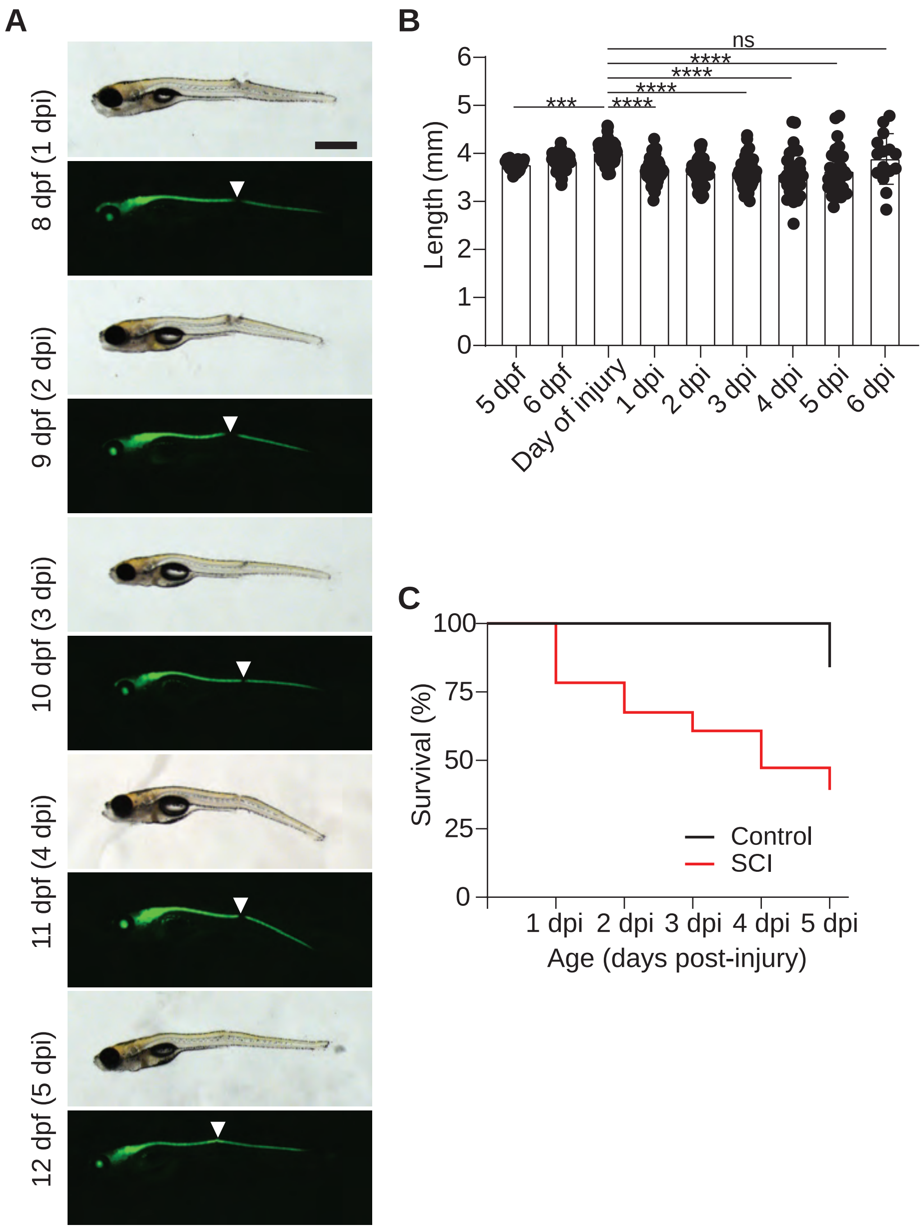

**Supplemental Figure 1. Rotifer-fed larvae have continued growth and regeneration after SCI** (A) Representative bright field and fluorescent micrographs of zebrafish larvae from 7 days post fertilization (dpf) through 13 dpf are shown. (B) Quantification of larval zebrafish length. Larvae were measured in Fiji and the average length of the larvae ± s.d. plotted. Values for individual fish are shown. The nonparametric Kruskal-Wallis test was used to determine significance. (C) Kaplan-Meier curves for control (black) and SCI (red). For all micrographs, larvae orientation is lateral view, anterior left. Scale bar: 1 mm. ***$p \leq 0.001$, ****$p \leq 0.0001$

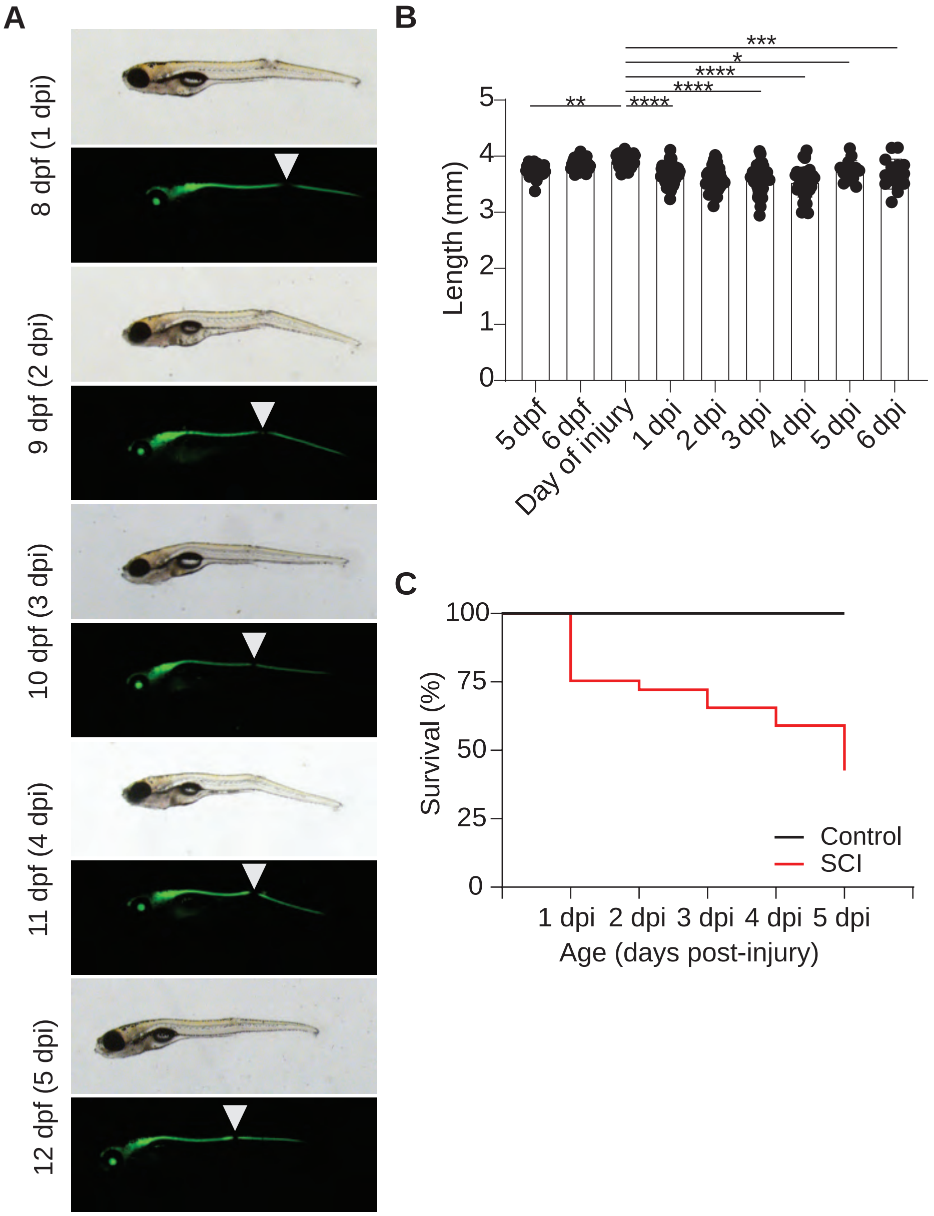

**Supplemental Figure 2. Powder-fed larvae have reduced growth and survival after SCI** (A) Representative bright field and fluorescent micrographs of zebrafish larvae from 7 days post fertilization (dpf) through 13 dpf are shown. (B) Quantification of larval zebrafish length. Larvae were measured in Fiji and the average length of the larvae ± s.d. plotted. Values for individual fish are shown. The nonparametric Kruskal-Wallis test was used to determine significance. (C) Kaplan-Meier curves for control (black) and SCI (red). For all micrographs, larvae orientation is lateral view, anterior left. Scale bar: 1 mm. *p ≤0.05, **p≤0.01, ***p≤0.001, ****p≤0.0001

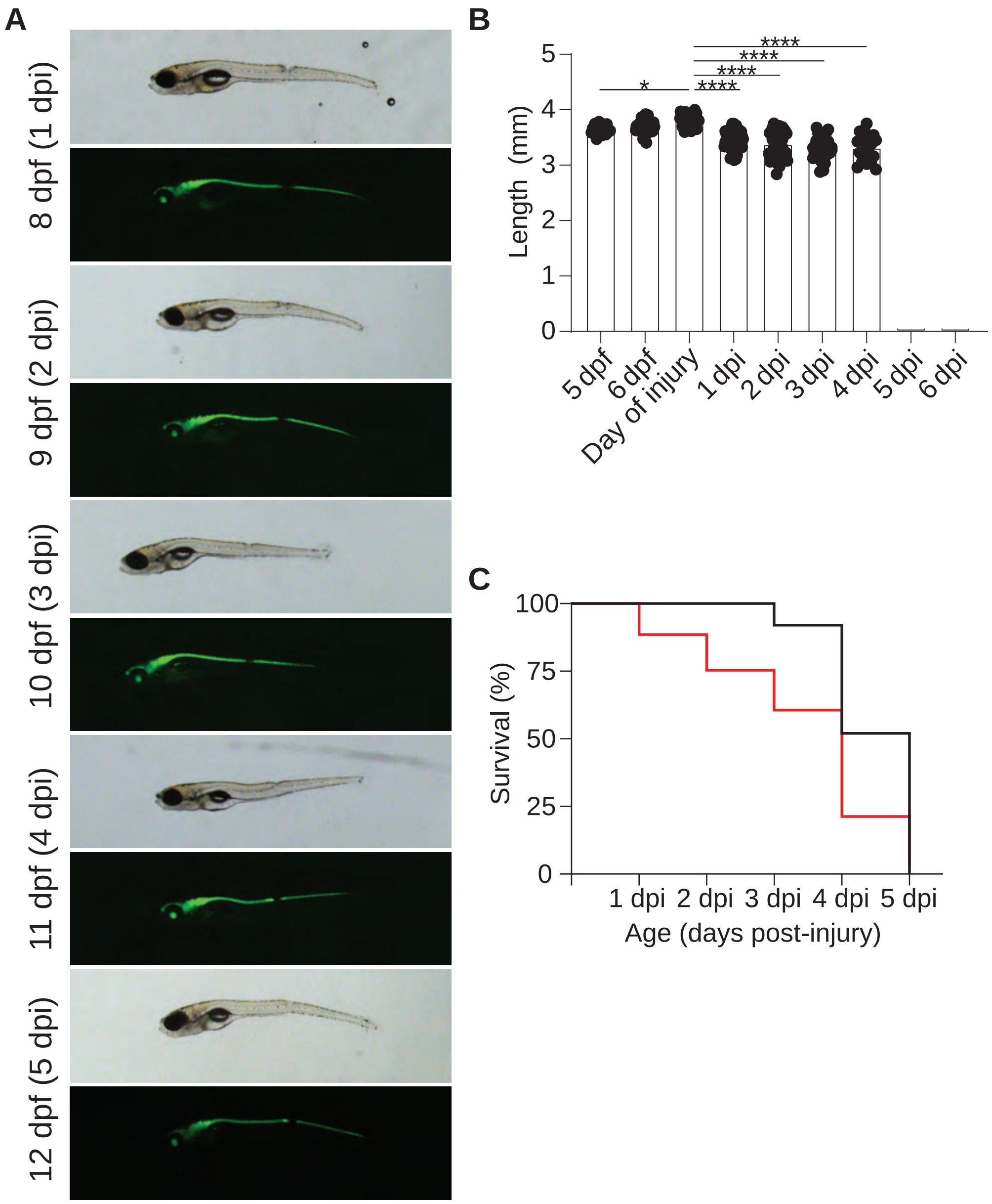

**Supplemental Figure 3. Nutrititve media cannot sustain life in older larvae** (A) Representative bright field and fluorescent micrographs of zebrafish larvae from 7 days post fertilization (dpf) through 12 dpf are shown. (B) Quantification of larval zebrafish length. Larvae were measured in Fiji and the average length of the larvae ± s.d. plotted. Values for individual fish are shown. The nonparametric Kruskal-Wallis test was used to determine significance. (C) Kaplan-Meier curves for control (black) and SCI (red). For all micrographs, larvae orientation is lateral view, anterior left. Scale bar: 1 mm. *p≤0.05, ****p≤0.0001
